# Familiarity training enhance straightening of neural trajectory for video prediction

**DOI:** 10.1101/2023.11.29.569215

**Authors:** Wentao Qiu, Ge Huang, Mirudhula Mukundan, Tai Sing Lee

## Abstract

Predictive processing in the visual system is pivotal for efficient sensory-driven behaviors. Previous research has shown that the visual system transforms sequential inputs into straighter temporal trajectories. However, the specific role of ‘neural straightening’ in predictive processing, especially in how learning reshapes this phenomenon for enhanced prediction, remains unclear. To address this, we analyzed V2 population activity in macaques during familiarity training with video stimuli. Our findings reveal that repeated exposure to the same movies significantly enhances neural straightening, indicating a critical role of learning in refining neural trajectories for prediction. In parallel, our studies with the deep predictive network model, PredNet, demonstrated similar enhancements in neural straightening in response to familiar movies. This underscores a strong association between neural straightening and predictive coding. Together, our results provide novel insights into the adaptive mechanisms of the visual cortex, enriching our understanding of how learning shapes neural pathways for efficient prediction.

## Introduction

The visual system’s capacity for predictive processing, essential for adaptive behavior in dynamic environments, hinges on its ability to interpret and anticipate sensory inputs. A recent advancement in understanding this capacity is the concept of ‘neural straightening’ (Hénaff et al., 2019, 2021), where the brain’s visual system is thought to refine neural trajectories in response to visual stimuli, thereby streamlining representational space to enhance prediction (Chung and Abbott, 2021). Despite its widespread recognition, direct evidence elucidating the functionality and mechanisms of neural straightening has been elusive.

This study addresses a fundamental question: How does neural straightening facilitate prediction, and what mechanisms underlie this process? We investigate the influence of experience and familiarity on neural straightening, employing both biological and computational models. In the biological domain, we analyze the V2 region population activity in macaques exposed to familiar and novel visual stimuli, exploring how experience alters neural trajectory straightness.

In parallel, our computational exploration centers on predictive coding within deep predictive networks, specifically PredNet. This model, grounded in predictive coding principles, uses error-driven learning to refine neural representations, offering a computational lens to examine neural straightening. Our novel approach in PredNet incorporates controlled noise, simulating learning dynamics akin to those in biological brains and revealing the intricate relationship between predictive coding and neural straightening.

Our findings in PredNet, where predictive coding not only supports temporal prediction but also critically influences neural straightening, mirror our biological observations. This synergy between biological and artificial systems in our study provides a comprehensive understanding of the role ofneural straightening in sensory prediction. By bridging biological perception and computational modeling, our research advances the knowledge of sensory processing, highlighting the impact of past experiences on the predictive abilities of neural systems. It paves the way for future explorations into the adaptability and cognitive functions of both biological and artificial visual systems.

## Results

### Enhanced Neural Straightening in Macaque Visual Cortex

Familiarity training is a well established approach to understand the neural plasticity and learning mechanisms in the primary visual cortex (Huang et al., 2018; Koyano et al., 2023; Price et al., 2023). In exploring the impact of familiarity training on neural trajectories, our study focused on two macaques, Gabby and Leo (Figure 1). They were exposed to both familiar and novel movie stimuli over several days—Gabby for 6 days and Leo across two experiments, one spanning 14 days and the other 11 days. We designated the initial and final phases of these sessions as ‘early’ and ‘late’ stages, respectively, to assess changes over time.

**Figure 1.**
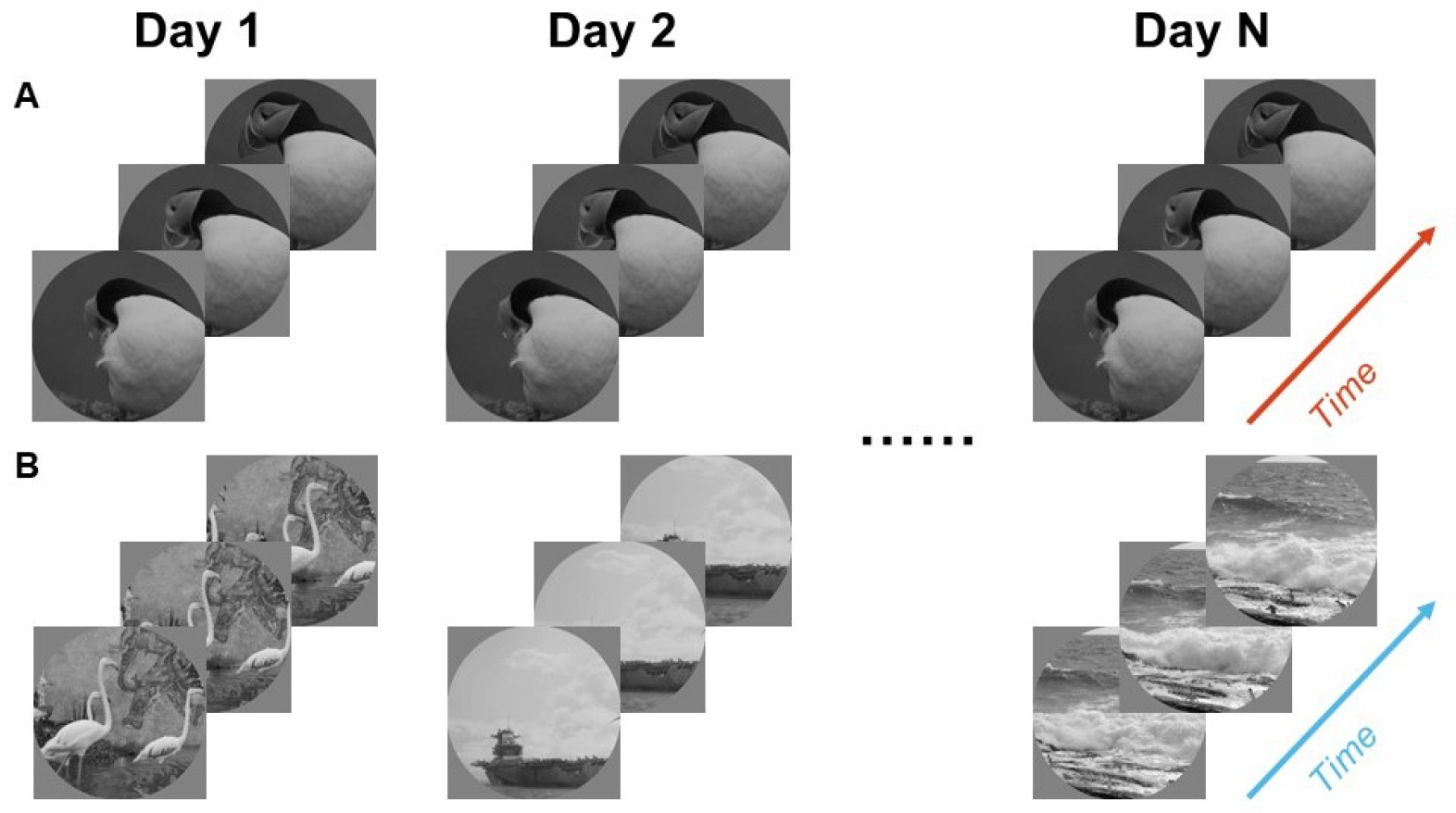
Familiarity training experiment. (**A**) The same familiar movies are repeatedly shown on each day. **B**) Different novel movies are shown each day.

Peristimulus time histogram (PSTH) analysis of the same electrodes on the first and last days (as illustrated in Figures 2, 3, and 4) revealed notable changes. Consistent with previous findings of familiarity suppression in V2 for familiar stimuli (Huang et al., 2018), we observed reduced neural responses to familiar movies in the late stage, whereas responses to novel movies re-mained largely unchanged from the early stage. Importantly, neural trajectories for familiar movies exhibited increased straightness over time.

**Figure 2.**
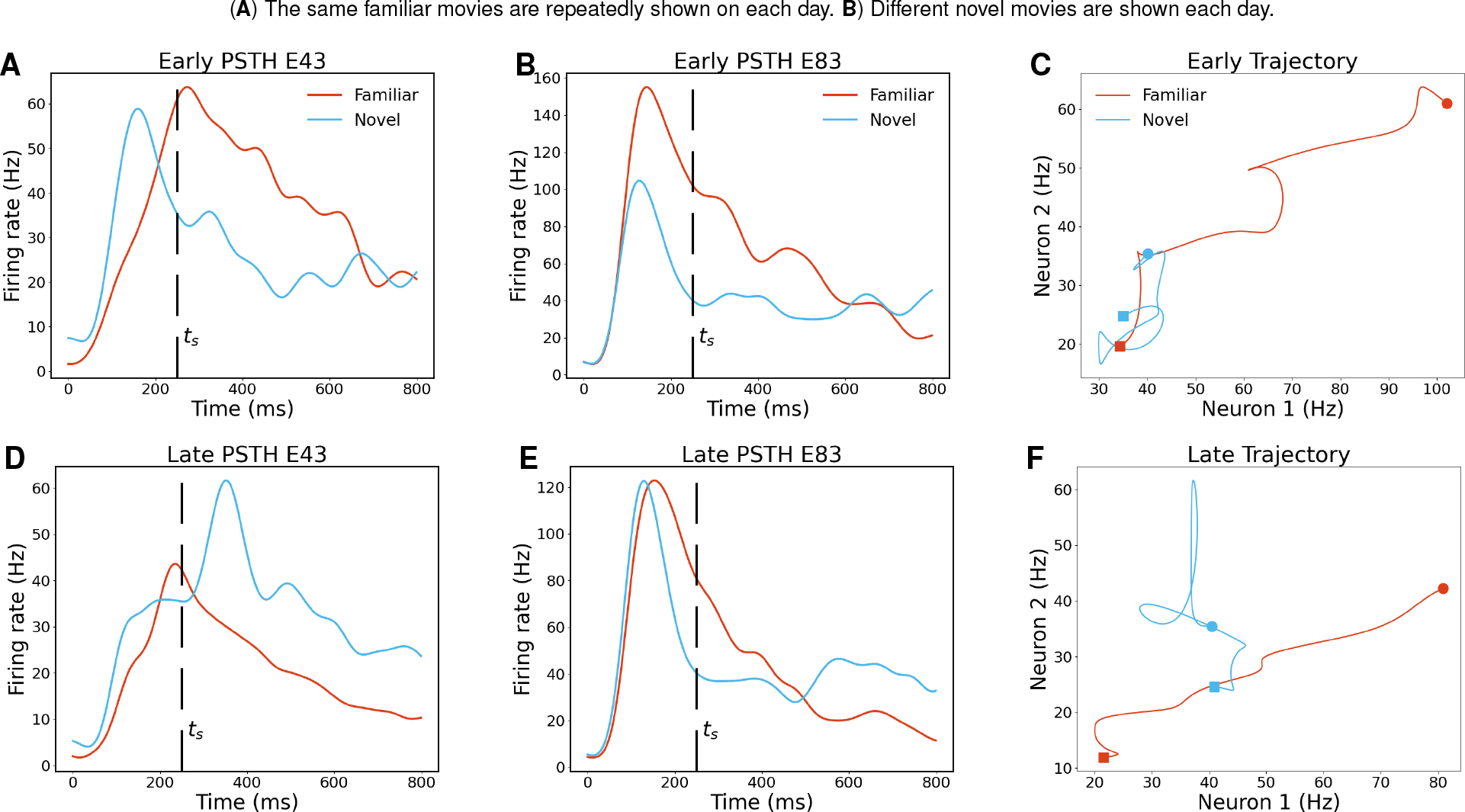
Illustration of reshaped neural trajectories in Gabby 1. (**A**) PSTH of Electrode 43 for one familiar and novel movie on the first day. **B**) PSTH of Electrode 83 for one familiar and novel movie on the first day. **C**) Two dimension neural trajectory of Electrode 43 and 83 for one familiar and novel movie on the first day. **D**) PSTH of Electrode 43 for one familiar and novel movie on the last day. **E**) PSTH of Electrode 83 for one familiar and novel movie on the last day. **F**) Two dimension neural trajectory of Electrode 43 and 83 for one familiar and novel movie on the last day.

**Figure 3.**
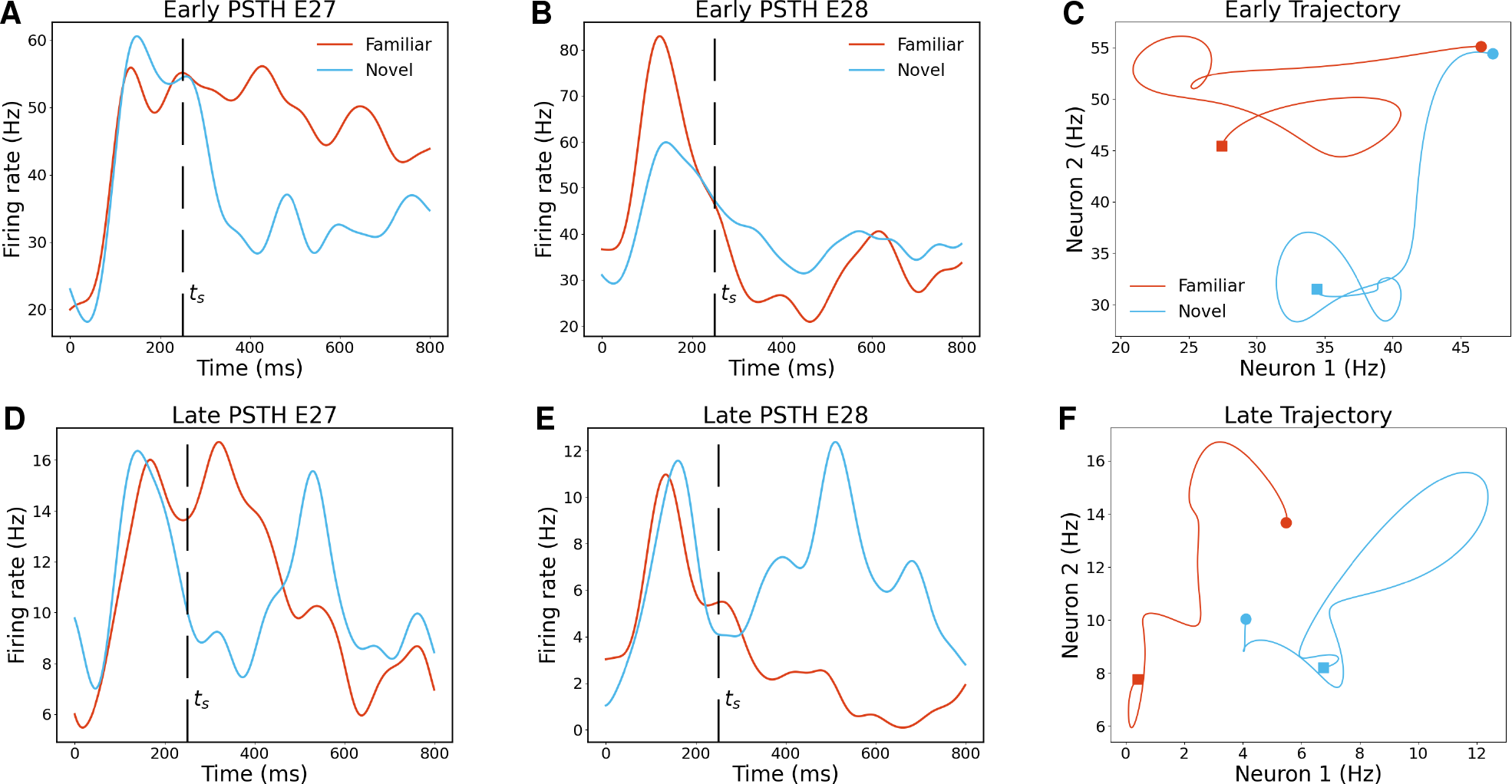
Illustration of reshaped neural trajectories in Leo 1. (**A**) PSTH of Electrode 27 for one familiar and novel movie on the first day. **B**) PSTH of Electrode 28 for one familiar and novel movie on the first day. **C**) Two dimension neural trajectory of Electrode 27 and 28 for one familiar and novel movie on the first day. **D**) PSTH of Electrode 27 for one familiar and novel movie on the last day. **E**) PSTH of Electrode 28 for one familiar and novel movie on the last day. **F**) Two dimension neural trajectory of Electrode 27 and 28 for one familiar and novel movie on the last day.

**Figure 4.**
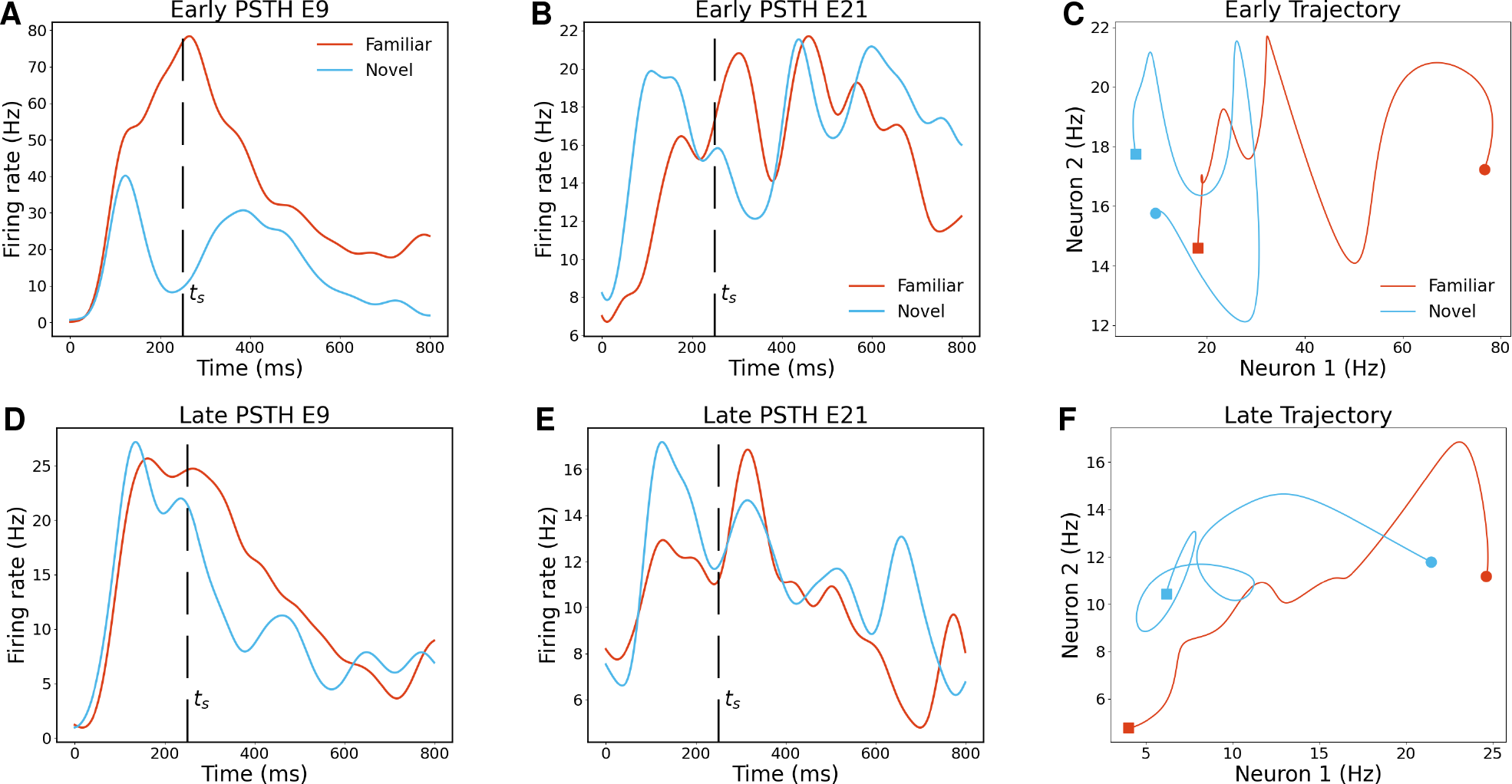
Illustration of reshaped neural trajectories in Leo 3. (**A**) PSTH of Electrode 9 for one familiar and novel movie on the first day. **B**) PSTH of Electrode 21 for one familiar and novel movie on the first day. **C**) Two dimension neural trajectory of Electrode 9 and 21 for one familiar and novel movie on the first day. **D**) PSTH of Electrode 9 for one familiar and novel movie on the last day. **E**) PSTH of Electrode 21 for one familiar and novel movie on the last day. **F**) Two dimension neural trajectory of Electrode 9 and 21 for one familiar and novel movie on the last day.

**Figure 5.**
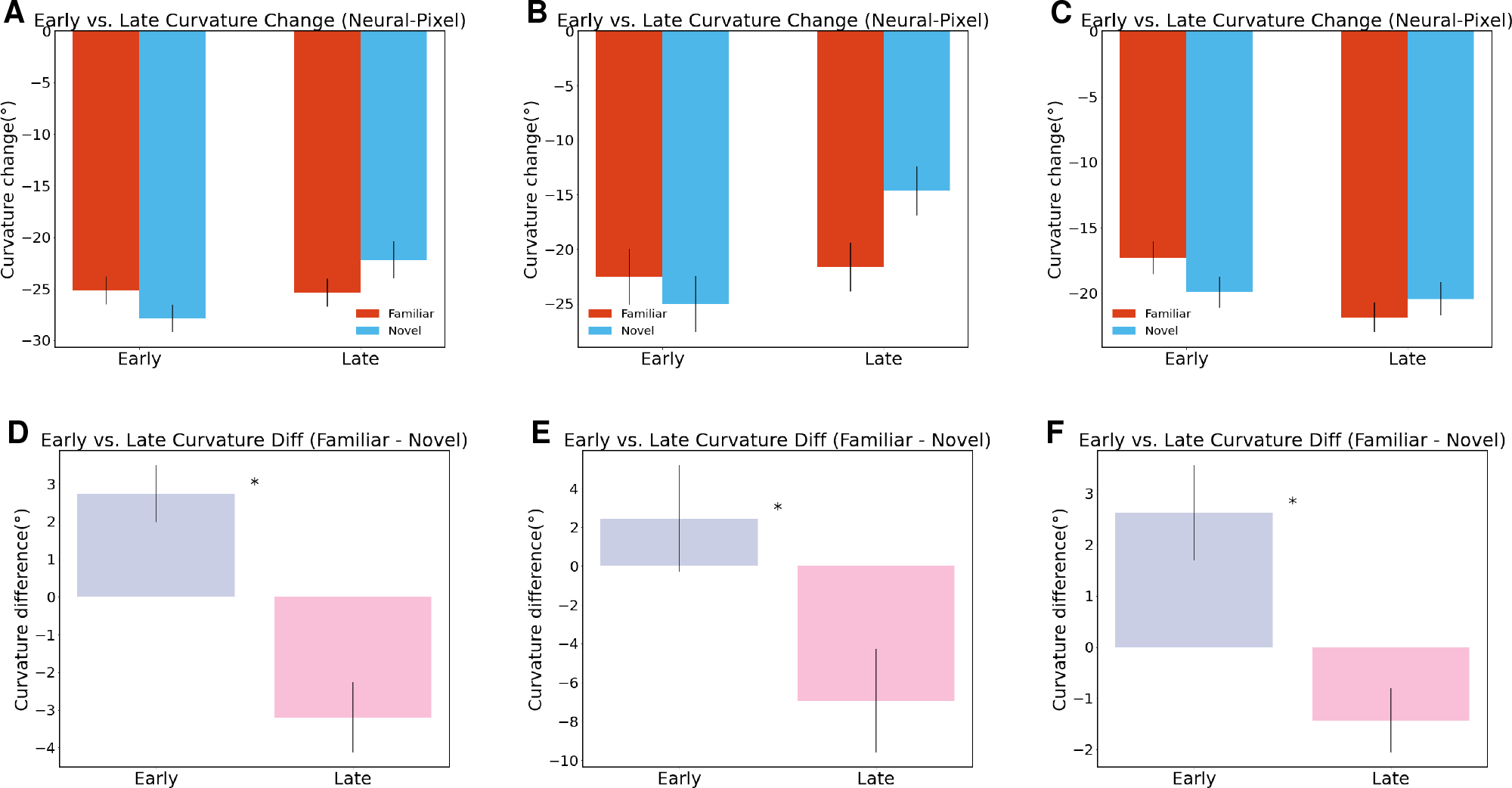
Familiarity training effects on neural straightening in macaque V2 cortex. (**A**) Changing neural straightening for Gabby 1. **B**) Changing neural straightening for Leo 1. **C**) Changing neural straightening for Leo 3. **D**) Neural straightening is enhanced at the late stage for Gabby 1. (independent t-test, p = 0.022) **E**) Neural straightening is enhanced at the late stage for Leo 1. (independent t-test, p = 0.025) **F**) Neural straightening is enhanced at the late stage for Leo 3. (independent t-test, p = 0.045)

**Figure 6.**
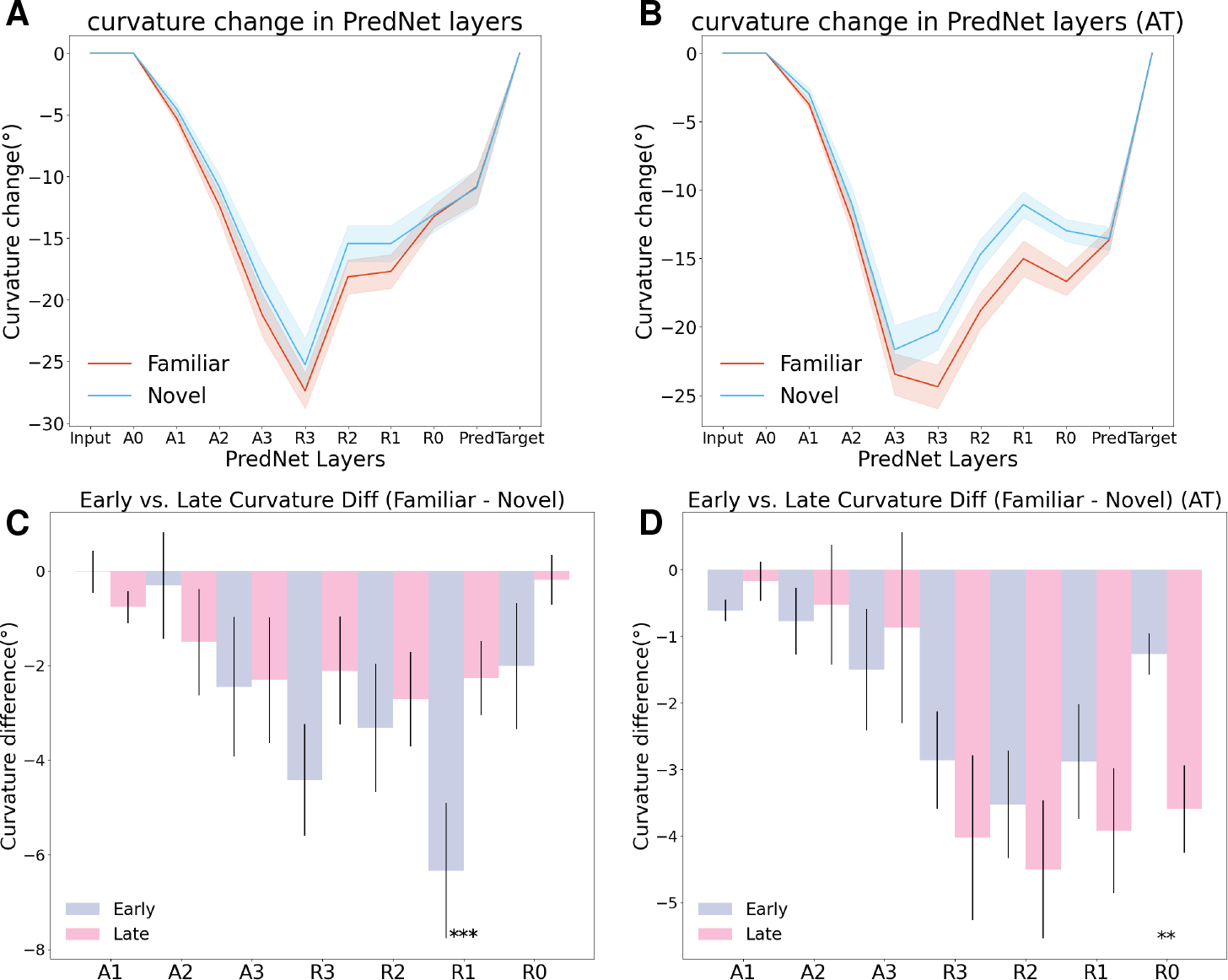
Familiarity training effects on neural straightening in PredNet. (**A**) Neural straightening for familiar and novel movies in standard training PredNet. **B**) Neural straightening for familiar and novel movies in adversarial training PredNet. **C**) Learning process of neural straightening in standard training PredNet. **D**) Learning process of neural straightening in adversarial training PredNet.

To quantify the trajectory straightening, we employed a short-time drift Gaussian Process Factor Analysis (STD-GPFA), as detailed in Section Methods. This method, an advancement over traditional GPFA, effectively captures non-linearities in latent neural dynamics. Using curvature measurements of the latent trajectories derived from neural spike counts, we observed significant decreases in curvature from early to late stages, indicating enhanced straightening (Gabby 1 *c*_*late*_ *− c*_*early*_ = *−*5.94^*°*^; Leo 1 *c*_*late*_ *− c*_*early*_ = *− −* 9.40^*°*^; Leo 3 *c*_*late*_ *− c*_*early*_ = *−*4.05^*°*^).

### Enhanced Neural trajectories in Deep Predictive Coding Networks for Video Prediction Tasks

Our investigation into PredNet, as proposed by (Lotter et al., 2016), centered around its application in frame prediction tasks using predictive coding, a principle fundamental to this model (Rao and Ballard, 1999). We aligned the training and test datasets in PredNet with familiar and novel movies, respectively, utilizing the KITTI dataset for training. We defined the early stage of the model after 10 epochs of training and the late stage after 100 epochs.

In our analysis, we discovered that PredNet, under standard training conditions, not only exhibited the neural straightening effect but also differentiated between familiar and novel movies, which is new compared with previous studies on straightening in PredNet (Harrington et al., 2022). Here, straightening is not forced as a objective function in the neural network, which is a common approach used in previous deep neural network models for the frame prediction tasks (Goroshin et al., 2015). This outcome aligns with our biological observations, indicating that predictive coding inherently generates neural straightening. However, a deviation was noted in the learning trajectory of PredNet compared to that of the macaque visual cortex, with a reduction in the straightening effect observed over the course of training, as shown in (Figure 6).

A pivotal finding emerged when we employed adversarial training using the Fast Gradient Sign Method (FGSM). This modification not only enhanced the neural straightening effect through training but also optimized PredNet’s predictive capabilities. This is consistent with previous research that adversarial learning mimic the neural representation in the biological systems ((Toosi and Issa, 2023; Guo et al., 2022). These results emphasize that neural straightening, fostered by predictive coding, is essential for efficient prediction in the visual system. Furthermore, they suggest that with appropriate training adaptations, PredNet can more accurately replicate the learning dynamics and prediction facilitation mechanisms inherent in biological systems.

We believe that this integrated approach in PredNet underscores the critical role of neural straightening within the framework of predictive coding. It highlights its importance in enhancing the visual system’s predictive accuracy, offering profound insights into the intersection of biological sensory processing and computational modeling.

## Discussion

Our study advances the understanding of ‘neural straightening’ in visual processing, building upon the foundational hypothesis that the visual system streamlines neural trajectories to enhance predictive accuracy. In our research, we have shown that this phenomenon is not only a passive outcome of visual processing but also an active component shaped by learning and experience, as evidenced in both macaque visual cortex and predictive coding networks.

Our findings in the macaque V2 region reveal that familiarity with visual stimuli significantly enhances neural straightening. This observation suggests an adaptive mechanism where the brain refines neural pathways to facilitate efficient prediction. It aligns with the notion that the brain’s processing of sensory information is not static but dynamically adjusts based on experience, an essential trait for navigating ever-changing environments.

In parallel, our exploration with PredNet, a deep predictive network model, reinforces the critical role of predictive coding in this process. We observed that predictive coding inherently generates neural straightening and enhances the model’s predictive capabilities, especially when subjected to adversarial training. This alignment with biological data suggests that neural straightening, fostered by predictive coding, is vital for efficient prediction in visual systems, both biological and artificial.

The significance of our work extends beyond the specific findings. It contributes to a broader understanding of how neural systems, whether biological or computational, use past experiences to optimize future predictions. This insight opens new avenues for developing more biologically plausible models of visual processing. Current deep neural networks, while effective in certain aspects of high-level visual function, often lack the nuanced capabilities inherent in biological systems, such as the ability to dynamically adapt to new information or to generalize from limited data. Incorporating principles like neural straightening and predictive coding could bridge this gap, leading to models that more accurately reflect the efficiency, flexibility, and robustness of biological intelligence.

Moreover, our study underscores the importance of a dual approach, combining biological and computational methods, to unravel the complex mechanisms of sensory processing. Just as the study of neural straightening in macaques sheds light on fundamental aspects of visual processing, so too does the examination of these principles in computational models. Together, they provide a more holistic view, one that is essential for advancing our understanding of both the brain and artificial intelligence.

In conclusion, our research not only provides empirical support for the role of neural straightening in sensory prediction but also highlights its potential as a guiding principle in the development of more advanced, biologically inspired computational models. This integration of biological insights with computational innovation promises to propel our understanding of sensory processing and cognition into new realms.

## Methods

### Measuring pixel-domain curvature

Given an image sequence, we can compare its curvature in the domain of pixel-intensities with that of neural responses. Measuring the pixel domain curvature is straightforward. Let 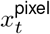 be the vector of pixel intensities of the frame at time *t ∈ {*0,…, *T }*. We define a sequence of normalized displacement vectors 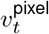

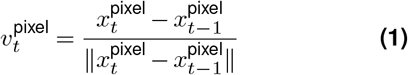

and the curvature at time *t* is simply the angle between two such vectors, which can be computed from their dot product:

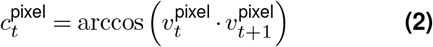

The global curvature of the sequence *c*_pixel_ is the average of these local curvature values, in degrees.

### Measuring neural-domain curvature in PredNet

Besides the curvatures in the pixel domain, we also calculate the neural curvatures in the PredNet in the same way, since the neural responses in PredNet don’t have trial variability.

### Measuring neural-domain curvature in Macaques Mathematical description of the drift GPFA model

We improve the Gaussian process factor analysis (GPFA) (Yu et al., 2008) model to measure the neural curvature. Let **y**_*t*_ *∈* ℝ^*q×*1^ be the high-dimensional vector of square-rooted spike counts recorded at time point *t* = 1, …, *T*, where *q* is the number of neurons being recorded simultaneously. We seek to extract a corresponding low-dimensional latent *neural state* **x**_*t*_ *∈* ℝ^*p×*1^ at each time point, where *p* is the dimensionality of the state space (*p < q*). For notational convenience, we group the neural states from all time points into a *neural trajectory* denoted by the matrix *X* = [**x**_1_, …, **x**_*T*_ ] *∈* ℝ^*p×T*^. Similarly, the observations can be grouped into a matrix *Y* = [**y**_1_, …, **y**_*T*_ ] *∈* ℝ^*q×T*^. We define a linear-Gaussian relationship between the observations **y**_*t*_ and neural states **x**_*t*_,

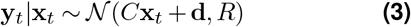

where *C ∈* ℝ^*q×p*^, **d** *∈* ℝ^*q×*1^, and *R ∈* ℝ^*q×q*^ are model parameters to be learned. As in FA, we constrain the covariance matrix *R* to be diagonal.

Originally, a separate GP is defined for each dimension of the state space indexed by *i* = 1, …, *p*,

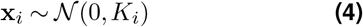

where **x**_*i*_ *∈* ℝ^1*×T*^ is the ith row of *X* and *K*_*i*_ *∈* ℝ^*T ×T*^ is the covariance matrix for the i-th GP.

The original model is not proper for measuring neural curvatures, since it’s assuming neural dynamics is doing pure GP, which will easy to give a 90^*°*^estimation (Consider an isotropic Motion, direction range is [0^*°*^,180^*°*^], the statistical mean is 90^*°*^). Therefore, we modify the latent neural dynamics by considering a more general version, called short-time drift GPFA model. We consider there’s a time-various speed at each time *V* = [**v**_1_, …, **v**_*T*_ ] *∈* ℝ^*p×T*^. At each time, the speed of each demension is indexed by *i* = 1, …, *p*,

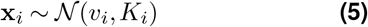

As the original model, we use squared exponential covariance function

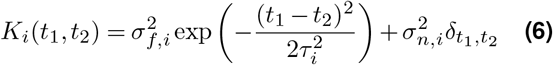

where *K*_*i*_(*t*_1_, *t*_2_) denotes the (*t*_1_, *t*_2_)th entry of *K*_*i*_ and *t*_1_, *t*_2_ = 1, …, *T*. The SE covariance is defined by its signal variance 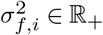, characteristic timescale *τ*_*i*_ *∈* ℝ_+_, and GP noise variance 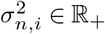 The Kronecker delta *δ*_*t*1_,_*t* 2_ equals 1 if *t*_1_ = *t*_2_ and 0 otherwise. The SE is an example of a stationary covariance; by modeling drift speed, we are able to catch other stationary and non-stationary covariances. (For example, illumination change is common in visual stimuli, which arouse a strong population activity at the beginning.)

### Fitting the short-time drift GPFA model

Since the time-various drift speed makes it hard to use traditional Expectation Maximization (EM) algorithm to optimize the model. We take advantage of the benefits from deep learning. We seek the model parameters ***θ*** = {*C, d, R, τ*_1_, …, *τ*_*p*_, *V*} that maximize the probability of the observed data **Y**. L2 loss is used for back-propagation to learn the parameters.

### Calculate the neural curvature

Since a time-various speed at each time *V* = [**v**_1_, …, **v**_*T*_ ] *∈* ℝ^*p×T*^ is inferred, we can directly use the speed to get the neural curvature with

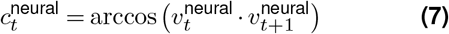

### Comparison with other methods

In our study, measuring neural straightening in the macaque visual cortex presented unique methodological challenges. Initially, one might consider linear or nonlinear dimension reduction methods, such as PCA or MDS, as direct approaches to extract latent neural trajectories. However, these methods proved insufficient, echoing limitations noted in previous research (Hénaff et al., 2021). Specifically, we found that neurons with high and fluctuating firing rates tend to dominate the principal components, biasing curvature measurements towards higher degrees. Additionally, the static linear dynamics model used in prior studies (Hénaff et al., 2021) was not suitable for our dataset, which differed significantly in stimulus presentation. Unlike the image sequence task where each image is followed by a period of constant luminance, our experiment involved continuous video presentation, introducing complex and nonlinear latent dynamics due to factors like gradual luminance changes post-stimulus onset and recurrent neuronal connections.

To address these challenges, we developed this novel model: the short-time drift Gaussian Process Factor Analysis (STD-GPFA). This model operates under the assumption that latent dynamics are linear within each adjacent frame of the video. This approach allowed us to effectively capture and quantify the neural straightening phenomenon and observe the impact of learning on this process. Our adaptation of GPFA to accommodate the temporal complexities of video stimuli represents a significant methodological advancement, offering a more nuanced understanding of neural trajectory dynamics in response to more dynamic and ecologically valid visual stimuli.

---

**Figure S1.**
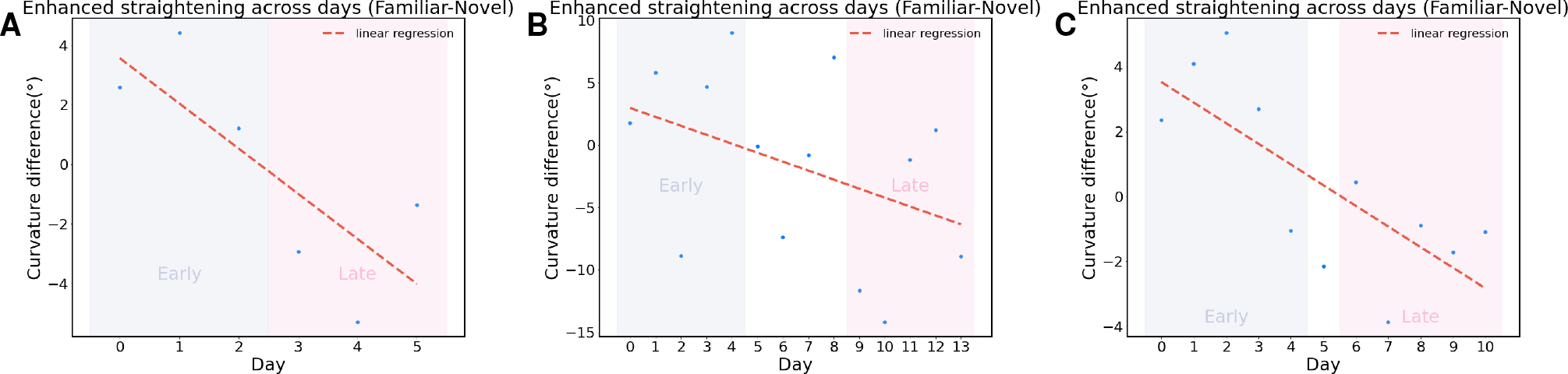
Familiarity training enhances neural straightening across days. (**A**) Changing neural straightening each day for Gabby 1. *slope* = *−*1.517, *r* = *−*0.781, *p* = 0.066 (**B**) Changing neural straightening each day for Leo 1. *slope* = *−*0.718, *r* = *−*0.409, *p* = 0.146 **C**) Changing neural straightening each day for Leo 3.*slope* = *−*0.636, *r* = *−*0.748, *p* = 0.008

## Supplementary Information

### A. Monkey detail

**Table 1.** Neuron and Electrode Counts in Leo Experiment 1

| Day | Neuron | Electrode | Day | Neuron | Electrode |
| --- | --- | --- | --- | --- | --- |
| 6/18/2015 | 13 | 7 | 6/30/2015 | 9 | 7 |
| 6/19/2015 | 11 | 7 | 7/02/2015 | 17 | 9 |
| 6/23/2015 | 8 | 7 | 7/03/2015 | 19 | 10 |
| 6/24/2015 | 12 | 7 | 7/04/2015 | 19 | 11 |
| 6/25/2015 | 13 | 8 | 7/07/2015 | 12 | 8 |
| 6/27/2015 | 14 | 8 | 7/08/2015 | 11 | 10 |
| 6/29/2015 | 13 | 8 | 7/09/2015 | 15 | 11 |

**Table 2.**
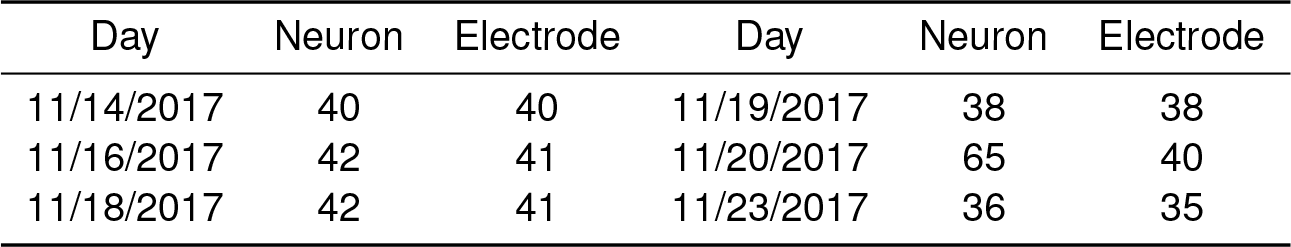
Neuron and Electrode Counts in Gabby experiment 1

| Day | Neuron | Electrode | Day | Neuron | Electrode |
| --- | --- | --- | --- | --- | --- |
| 11/14/2017 | 40 | 40 | 11/19/2017 | 38 | 38 |
| 11/16/2017 | 42 | 41 | 11/20/2017 | 65 | 40 |
| 11/18/2017 | 42 | 41 | 11/23/2017 | 36 | 35 |

**Table 3.** Neuron and Electrode Counts for Leo experiment 3

| Day | Neuron | Electrode | Day | Neuron | Electrode |
| --- | --- | --- | --- | --- | --- |
| 8/18/2015 | 31 | 19 | 8/28/2015 | 34 | 21 |
| 8/19/2015 | 35 | 23 | 8/31/2015 | 26 | 19 |
| 8/20/2015 | 34 | 21 | 9/01/2015 | 30 | 22 |
| 8/21/2015 | 29 | 19 | 9/02/2015 | 27 | 21 |
| 8/24/2015 | 30 | 20 | 9/03/2015 | 31 | 20 |
| 8/25/2015 | 27 | 18 |  |  |  |

